# Identification of Molecular Compounds Targeting Bacterial Propionate Metabolism with Topological Machine Learning

**DOI:** 10.1101/2024.10.26.620439

**Authors:** Astrit Tola, Shan Aziz, Dannie Zhabilov, Duane Winkler, Baris Coskunuzer, Mehmet Candas

**Affiliations:** University of Texas at Dallas, Dept. Math. Sciences, Richardson, TX 75080; University of Texas at Dallas, Dept. Biological Sciences, Richardson, TX 75080

**Keywords:** Antimicrobial development, topological data analysis, ligand-based virtual screening, methylcitrate pathway

## Abstract

This study demonstrates the effective integration of comparative protein sequence analysis with novel topological machine learning methods to tackle a key issue in computational biology: identifying potential inhibitor compounds for methylcitrate dehydratase—an enzyme essential to the methylcitrate pathway in bacteria and fungi. While many ML models have proven effective on benchmark datasets, we applied these techniques specifically to discover compounds for this target protein. This pathway, crucial for metabolizing propionic acid, is essential in pathogenic bacteria for utilizing host-derived lipids and amino acids, making it a promising antimicrobial target. Inefficient removal of propionate can lead to toxic accumulation, threatening microbial survival.

In our research, we utilize the latest methods in topological machine learning, *multiparameter persistence*, to transform the molecular structure of potential compounds into topological vectors for ligand-based drug discovery. By applying our tailored topological model, we prioritized 15 compounds with promising characteristics for inhibiting the active site of methylcitrate dehydratase. Computational molecular docking simulations reveal that these identified compounds interact with key amino acid residues critical to the enzyme’s function. Our findings underscore the power of integrating topology-based modeling with comparative sequencestructure-function analysis and ligand docking. Considering compounds’ geometric fit, energy, and interaction profiles enhances predictions and guides the design of optimized new compounds. This refined approach offers a solid foundation for lead optimization, representing the most promising molecular scaffolds for modification, advancing compound discovery, and providing valuable insights into binding interactions for further experimental validation and potential drug development. Our code is available at https://github.com/AstritTola/Molecular-Compounds-Targeting

## 1 Introduction

The landscape of drug discovery is undergoing a significant transformation with the integration of machine learning (ML) and virtual screening techniques. These advanced computational methods facilitate rapid in silico studies involving protein targets, enabling the identification and validation of potential drug leads with remarkable speed and accuracy. This acceleration not only expedites the drug discovery process but also offers valuable insights into the interaction modalities between lead compounds and target proteins [35]. As a result, the synergy between ML and virtual screening revolutionizes drug development, leading to the creation of effective therapeutic agents within considerably shortened timeframes [36].

The urgent rise of antimicrobial resistance underscores the necessity for innovative treatment strategies [47,30,39]. Traditional antibiotics are increasingly becoming ineffective against resistant strains, highlighting the critical need to explore new avenues for antimicrobial development [11]. By focusing on previously unexploited bacterial proteins and biochemical pathways that lack mammalian counterparts, researchers can disrupt essential bacterial processes in novel ways. This approach not only expands the arsenal of antimicrobial agents but also enhances the chances of discovering treatments with unique modes of action. Targeting these underexplored bacterial pathways is crucial to combating antimicrobial resistance and ensuring the future efficacy of antimicrobial therapies [32].

One promising area of exploration is the utilization of propionic acid (or propionate), a three-carbon fatty acid that is prevalent in soil and serves as an intermediate in various metabolic pathways within cells. For example, it is produced during the oxidation of odd-chain fatty acids and several amino acids, as well as during cholesterol metabolism, where it forms bile acids. In mammals, propionyl-CoA, the active form of propionate, is converted into succinate through the methylmalonate pathway, involving biotin-dependent propionyl-CoA carboxylase and B12-dependent methylmalonyl-CoA mutase. In contrast, many bacteria and fungi primarily process propionate through the methylcitrate pathway, which converts propionyl-CoA into succinate and pyruvate [25,8,38]. The enzymes that facilitate the methylcitrate pathway in bacteria are encoded by the prp operon [18,20]. Although there are evolutionary variations in the structure and organization of this operon across different bacterial species, the pathway generally comprises four essential enzymatic reactions (see Figure 1). Initially, propionyl-CoA is synthesized from propionate by propionyl-CoA synthase (PrpE). This compound then condenses with oxaloacetate to form (2S,3S)-methylcitrate, catalyzed by methylcitrate synthase (PrpC). The subsequent stereospecific isomerization of (2S,3S)-methylcitrate to (2R,3S)-2-methylisocitrate is mediated by 2-methylcitrate dehydratase (AcnD), an enzyme with properties analogous to aconitase in the tricarboxylic acid (TCA) cycle. While AcnD is the primary enzyme for this reaction, additional bacterial proteins, such as PrpD and PrpF, may facilitate this process, suggesting that some variations may enhance metabolic robustness and environmental adaptability in some bacteria [34,16,6]. Finally, the enzyme PrpB cleaves (2R,3S)-2-methylisocitrate to yield pyruvate and succinate, which are integral to central metabolism and energy production. [6].

**Fig. 1:**
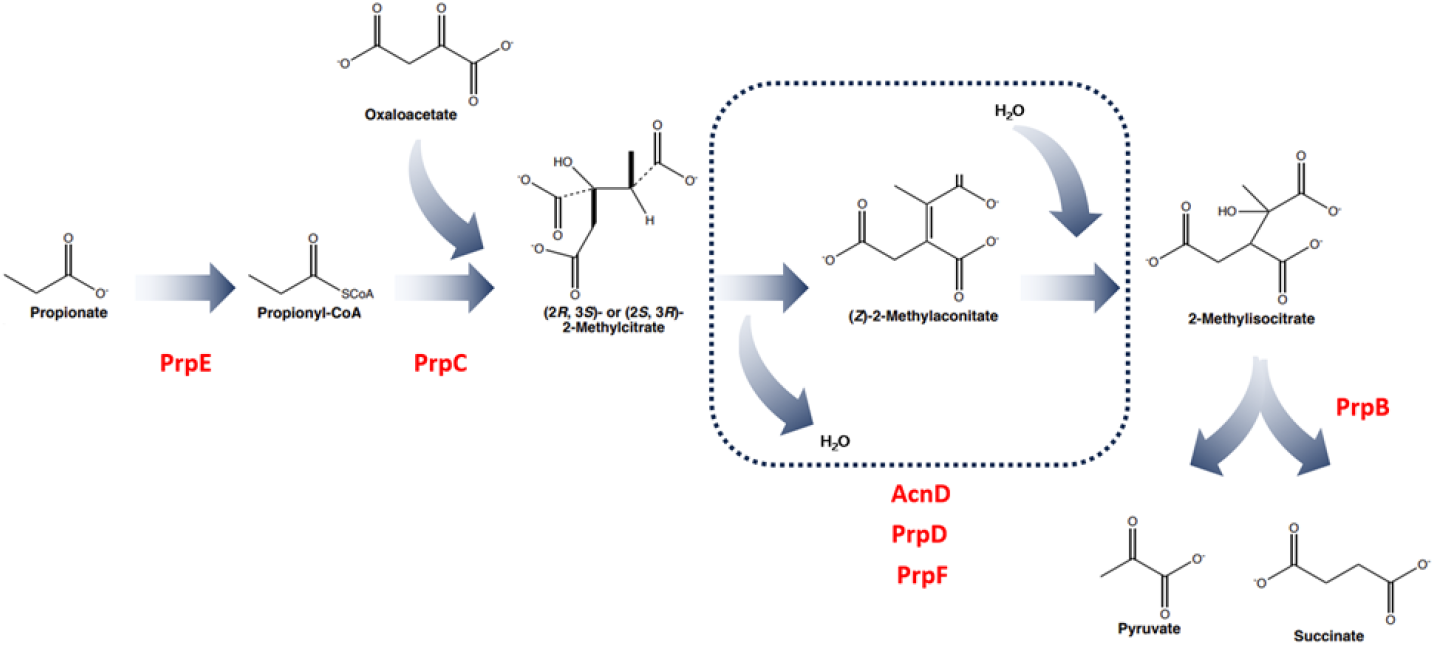
Methylcitrate pathway. The enzymes involved in the pathway are encoded by the prp operon in bacteria. While there are evolutionary variations in the structure and organization of the genes in the prp operon across different bacterial species, the methylcitrate pathway generally involves four essential enzymatic reactions: (1) formation of propionyl-CoA catalyzed by PrpE; (2) condensation of propionyl-CoA with oxaloacetate catalyzed by PrpC; (3) stereospecific isomerization of 2-methylcitrate to 2-methylisocitrate catalyzed by AcnD; and (4) cleavage of 2-methylisocitrate to form pyruvate and succinate by PrpB.

While propionate can serve as a carbon source, its accumulation can be toxic to bacteria and fungi, leading to growth suppression, even in glucose-rich environments. In this context, the methylcitrate pathway acts both as a catabolic and detoxification route [41].

The stereospecific isomerization of (2S,3S)-methylcitrate to (2R,3S)-2-methylisocitrate represents a critical intervention point in the methylcitrate pathway. Targeting this reaction with specific inhibitors offers a viable strategy to block propionate metabolism in bacteria. Such inhibition can lead to the accumulation of toxic intermediates, disrupting key metabolic processes and causing energy deficits and metabolic imbalances. Consequently, targeting AcnD emerges as a strategic focal point for antimicrobial drug development [41,14,7]. Significantly, AcnD is employed in the propionate metabolism of several gram-negative bacteria, including Shewanella, Vibrio, Neisseria, Pseudomonas, Acinetobacter, Ralstonia, and Burkholderia, which are associated with various diseases in humans, animals, and plants [17]. For instance, Pseudomonas aeruginosa is notorious for causing difficult-to-treat infections in clinical settings, especially in acute or chronic infections associated with immunocompromised patients, including individuals with chronic obstructive pulmonary disease (COPD), cystic fibrosis, cancer, traumas, burns, sepsis, and ventilator-associated pneumonia [31,33]. Since AcnD has no mammalian counterpart, inhibiting it provides a selective approach to disrupt propionate metabolism in pathogenic bacteria, controlling their growth and virulence [10].

In this study, we employed a novel approach to identify and characterize molecular compounds targeting AcnD to inhibit bacterial propionate catabolism. By utilizing three-dimensional structural information on protein-ligand interactions, we aimed to enhance rational drug discovery, despite the challenges posed by the dynamic nature of these interactions in identifying specific ligands. To tackle this, we adopted a ligand-based discovery approach, using mathematically modeled structural fingerprints complemented by an ML framework for iterative screening of virtual chemical libraries. Through this methodology, we identified and ranked 15 novel compounds targeting the active site of AcnD based on structural similarities with its native ligand, methylcitrate. Our docking experiments verify our results and show that these compounds can inhibit the methylcitrate pathway, thereby reducing bacterial growth and virulence, offering a promising new strategy to combat drug-resistant infections.

### 1.1 Related Work

#### Topological Methods for Drug Discovery

Topological methods have emerged as a vibrant area of research in computer-aided drug design in recent years, providing a powerful framework for understanding and predicting ligand-protein interactions by emphasizing the spatial arrangement and connectivity of atoms. In [24,12,29], the authors applied various forms of multiparameter persistence along with diverse vectorization techniques to develop highly effective topological fingerprinting methods for chemical compounds. Their methods demonstrated strong performance in benchmark datasets, excelling in both virtual screening and molecular property prediction tasks. Similarly, in [28], localized weighted persistent homology was introduced to capture local functional properties of biomolecules, proving effective in differentiating between DNA types. In [42], topological methods were integrated with graph neural networks to create the TopNet model, which achieved notable success in molecular property prediction and co-designing antibody sequences and structures using antigen-antibody complex datasets. While these approaches show significant promise as theoretical studies and perform well on benchmark datasets, there remains a gap in validating these models on real-world biological problems. In this study, we present a novel topological model for drug discovery targeting a specific protein and demonstrate its effectiveness through molecular docking experiments.

#### Antimicrobial Design

Computational and bioinformatics studies are crucial in gaining a deep understanding of metabolic and regulatory pathways and enzyme structures, which are essential for identifying potential targets for the development of new antimicrobials. Molecular structure-based approaches facilitate to design more effective strategies and inhibitors that specifically target critical proteins and biochemical reactions. For instance, by mapping out the exact structure of potential target enzymes, it is possible to predict how different molecules will interact with it, leading to the selection and/or creation of inhibitors that can block its function. Additionally, computational methods enable access to unexplored chemical space through customized virtual compound libraries. ML and generative models hold significant promise in this area for the identification of compounds capable of specifically interacting with target proteins [26,37].

Recent examples of computational biology in antimicrobial drug design include the use of ML-based classifiers for detecting antibacterial peptides and their targets [4,1,27]. This approach leverages large datasets and advanced algorithms to predict how different peptides interact with bacterial proteins, enabling the design of new antimicrobial agents [43]. Another example involves using genome-scale metabolic modeling to find new metabolic targets and virtual screening to discover selective enzyme inhibitors [22,45]. These examples highlight the power of combining computational tools with protein structure analysis to identify and prioritize tractable novel drug targets and develop effective drugs that can specifically and selectively disrupt metabolic enzymes. Overall, the identification of drug candidates from vast chemical libraries and predicting their binding specificity and affinity with molecular docking studies supports validation and optimization experiments, accelerating the drug discovery process and enhancing the precision and efficacy of new antimicrobial therapies, particularly against infections caused by resistant bacteria like Pseudomonas aeruginosa, which is increasingly becoming more difficult to treat [33,46].

## 2 Topological Methods for Compound Identification

In this section, we present our methodology. We begin by providing a brief overview of single persistent homology (PH), the foundational technique of our approach. Next, we introduce an extension of traditional PH, multiparameter persistence, incorporating multiple filtration functions from the domain to generate more robust and effective topological feature vectors for each compound. Finally, we combine these topological vectors with various ranking algorithms to identify the most promising compounds for our target (See Figure 2).

**Fig. 2:**
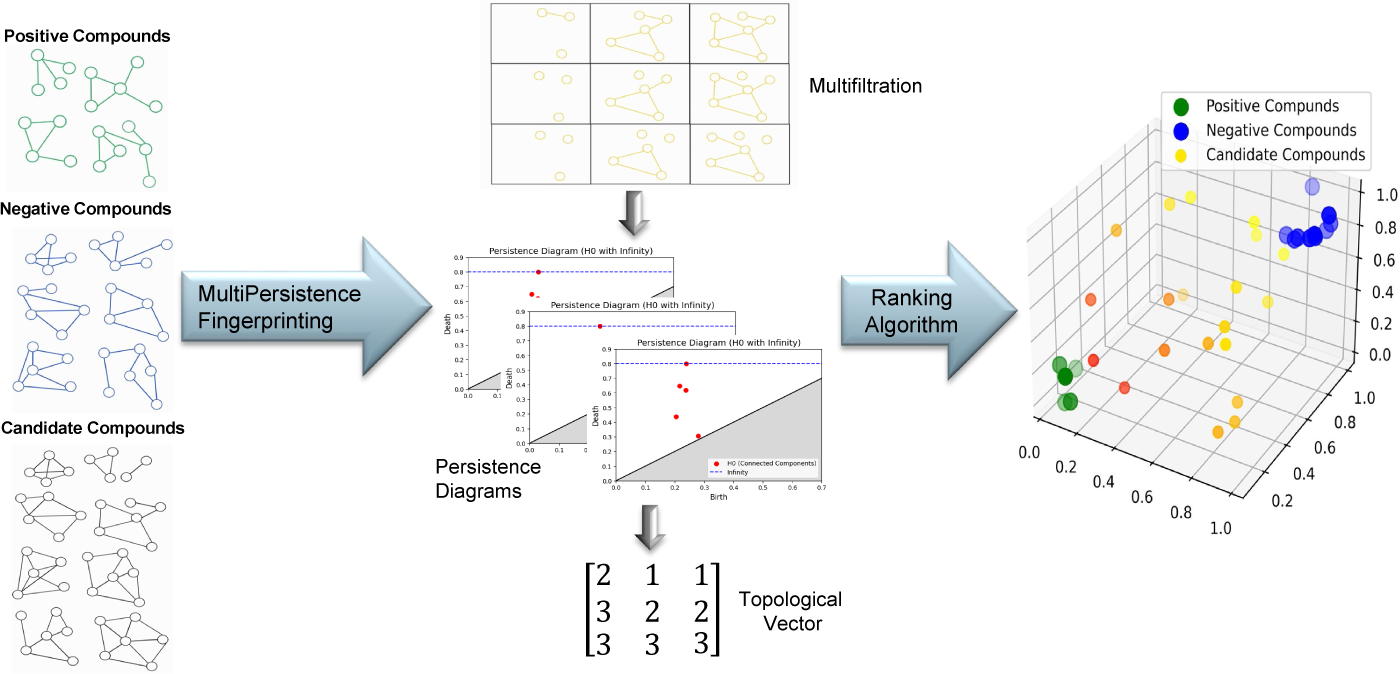
Topological Fingerprinting & Ranking. We first extract MultiPersistence vectors for each compound. Using these vectors, we fine-tune ranking algorithms with positive and negative compounds, allowing us to rank candidate compounds based on their topological representations.

**Fig. 3:**
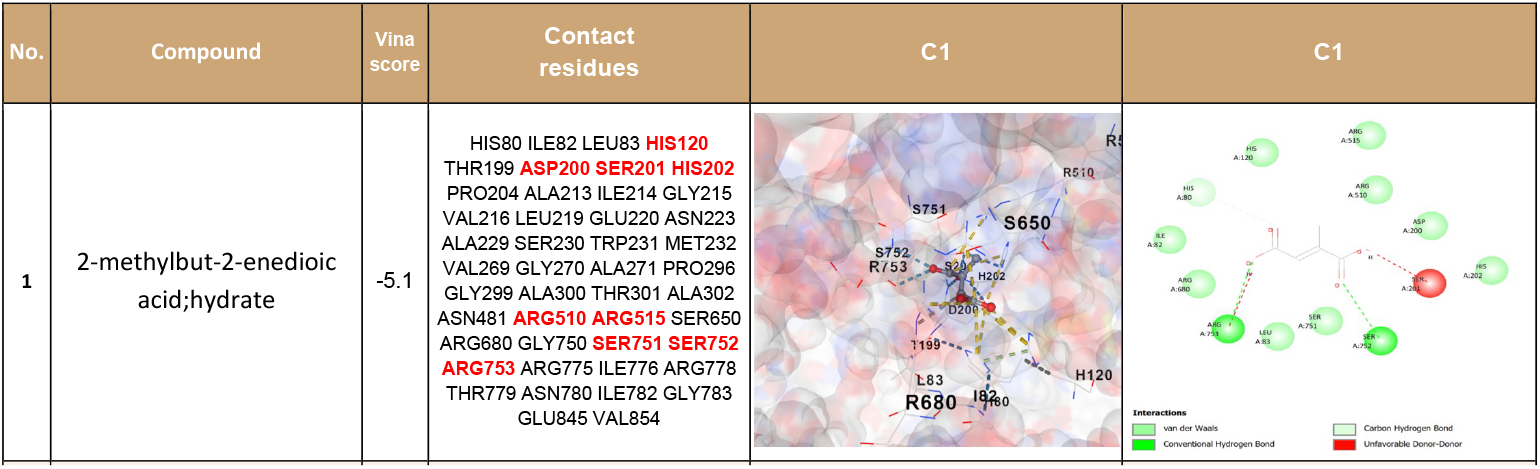
Docking simulations. We present a simulation result for one compound. Docking simulations for other 15 compounds are given in Figures 8 to 10 in Appendix.

**Fig. 4:**
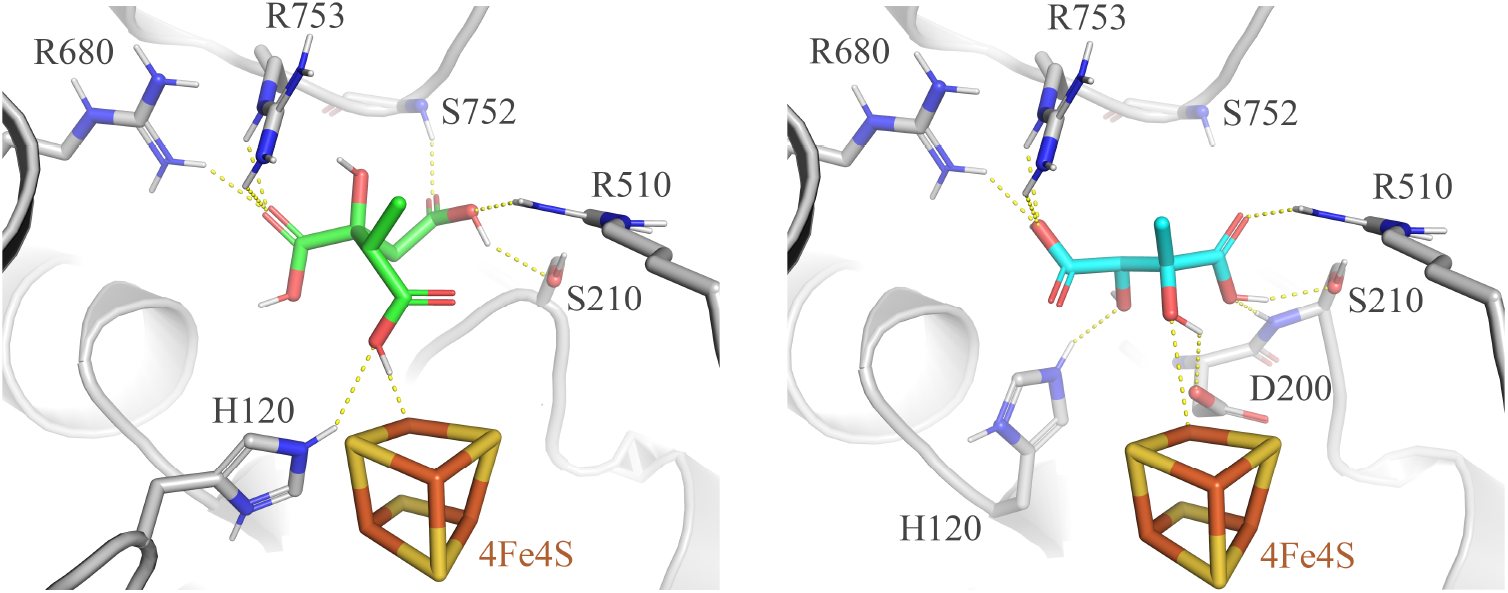
The left panel shows 2-methylcitrate (substrate - green) docked into the AcnD active site using the AutoDock Vina software suite. The AcnD model was generated using a combination of AlphaFold2 for the overall protein fold and structure-based manual placement of the 4Fe4S cluster using pig heart aconitase as an alignment protein. The substrate molecule binds within the active site similar to that of substrate molecules in related aconitase structures by utilizing a network of conserved arginine and serine residues. The right panel shows the docking of 2,3-dihydroxy-2-methylbutanedioic acid (cyan), which was identified in this study. The compound sits in a nearly identical orientation to the substrate and utilizes the same conserved active site residues as methylcitrate, in addition to another conserved residue (D200). The docking is obtained without providing initial details on active site location, size or critical liganding residues.

### Persistent Homology

To capture the deeper, often hidden structural properties of molecular graphs, we employ Persistent Homology (PH), a central tool in Topological Data Analysis (TDA). Unlike traditional methods that primarily focus on combinatorial or metric properties, PH allows us to quantify topological features—such as clusters, loops, and voids—that persist across multiple scales. By tracking the evolution and persistence of these features, PH unveils intricate patterns within the data that may otherwise remain undiscovered. This topological perspective is broadly applicable beyond graphs, extending to point clouds, images, and other complex datasets. For a comprehensive overview of PH’s application across different data domains, see [13,9].

The PH process can be summarized in three main steps. Let 𝒢 = (𝒱, ℰ) represent a graph with a node set 𝒱 and an edge set ℰ. The first step is *to construct a filtration*, a nested sequence of simplicial complexes derived from the graph. A typical approach involves constructing a nested sequence of subgraphs 𝒢_1_ ⊆ … ⊆ 𝒢_*N*_ = 𝒢, guided by a filtration function *f* : 𝒱 → ℝ and a set of thresholds ℐ = {*ϵ*_*i*_}, where *ϵ*_1_ = min_*v*∈𝒱_ *f* (*v*) *< ϵ*_2_ *<* … *< ϵ*_*N*_ = max_*v*∈𝒱_ *f* (*v*). For each *ϵ*_*i*_, a subset 𝒱_*i*_ = {*v*_*r*_ ∈ 𝒱 | *f* (*v*_*r*_) ≤ *ϵ*_*i*_} is formed, followed by the construction of the induced subgraph 𝒢_*i*_ = (𝒱_*i*_, ℰ_*i*_), where ℰ_*i*_ = {*e*_*rs*_ ∈ ℰ | *v*_*r*_, *v*_*s*_ ∈ ℰ_*i*_}. This step can be considered as decomposing the compound into subtle substructures in an organized way. Next, from each subgraph, we construct a simplicial complex, yielding a filtration 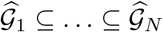. A common choice is to use clique complexes, where the *k*-simplices of a graph are formed by adding *k*-dimensional simplices for every complete (*k* + 1)-subgraph in 𝒢_*i*_ [2].

The second step is *to compute persistence diagrams*, where we track the evolution of topological features in the filtration 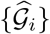. These features—such as connected components (0-holes), loops (1-holes), and cavities (2-holes)—are encoded by *k*-dimensional homology classes or *k*-holes. Using homology theory [19], PH captures the birth and death of each topological feature as we move through the filtration, recording these events in persistence diagrams. Specifically, the *k*-th persistence diagram PD_k_(𝒢) consists of pairs (*b*_*σ*_, *d*_*σ*_), where *b*_*σ*_ and *d*_*σ*_ represent the birth and death times of the *k*-hole *σ* in the filtration.

The final step is *vectorization*, which transforms persistence diagrams into a format suitable for ML applications. While PH provides detailed topological information as persistence diagrams, which is collections of points in ℝ^2^, they are not directly usable for most ML algorithms. To address this, a common technique is to vectorize the persistence diagrams into vectors [3]. These representations enable the smooth integration of topological signatures into ML workflows.

### Topological Vectors for Compounds

In the construction above, single persistent homology employs a single filtration function to obtain the sequence 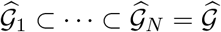. However, employing multiple filtration functions enables a more detailed analysis of the data. In particular, two node functions *f* : 𝒱 → ℝ and *g* : 𝒱 → ℝ, which provide complementary information about the network, can be combined to obtain a much finer decomposition of the data to generate a unique topological vector. In our case, these functions can be atomic number, bond strength (edge function), ionic energy, and other chemical functions. When paired, these functions induce a multifiltration functions *F* : 𝒱 → ℝ^2^ defined by *F* (*v*) = (*f* (*v*), *g*(*v*)) [9]. Next, we define sets of increasing thresholds 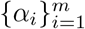 for *f* and 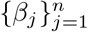 for *g*. Then, we have 𝒱_*ij*_ = {*v*_*r*_ ∈ 𝒱 | *f* (*v*_*r*_) ≤ *α*_*i*_, *g*(*v*_*r*_) ≤ *β*_*j*_}, which can be written as 𝒱_*ij*_ = *F*^−1^((−∞, *α*_*i*_] *×* (−∞, *β*_*j*_]). Let 𝒢_*ij*_ be the subgraph of 𝒢 induced by 𝒱_*ij*_, meaning the smallest subgraph of 𝒢 generated by 𝒱_*ij*_. This setup induces a bifiltration of complexes 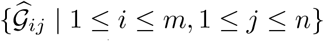, which can be visualized as a rectangular grid of size *m × n* (See Figure 6 in Appendix).

After obtaining multifiltration {𝒢_*ij*_}, we use *m* horizontal slices in the multipersistence module to obtain our topological vectors as follows. For each 1 ≤ *i*_0_ ≤ *m*, the row 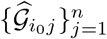 induces a single filtration and corresponding persistence diagram. The next step is to vectorize these *m* persistence diagrams. While one can use different vectorizations, e.g., Persistence Landscape, Silhouette, and Persistence Image, we employ Betti vectorization ***β*** for computational efficiency. Then, let 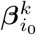 be the corresponding 1 *× n* Betti vector for 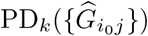. Now, we are ready to define our topological summary **M**_*φ*_ which is a 2*D*-vector (a matrix) 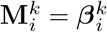 for 1 ≤ *i* ≤ *m*, and *k* = 0, 1 where 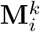 is the *i*^*th*^-row of **M**^*k*^. Hence, **M** is a 2*D*-vector of size *m × n* (See Appendix A.1 for explicit example).

In a way, we look at 𝒢 with a 2D lens with a finer resolution {𝒢_*ij*_} and we keep track of the evolution of topological features in the induced bifiltration 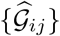. The main advantage of this technique is that the outputs are fixed-size matrices (or arrays) for each dataset which is highly suitable for various ML models.

## 3 Experiments and Results

### Compound Library

Inspired by contrastive learning [21], our library consists of three categories of compounds: *positive, negative*, and *candidate compounds*. Positive compounds are those known to bind to the target, labeled as *1*, and consist of 13 compounds. For the negative compound set, we generated artificial compounds by first identifying the type and number of non-hydrogen atoms in each positive compound. We then maintained the overall count of non-hydrogen atoms while altering the atom-type distribution (e.g., carbon, oxygen, nitrogen) to ensure a different composition from the original compound. At the end, 153 compounds were obtained. These compounds are labeled *0*.

In constructing the candidate compound database, we focused on four key compounds: *methylcitrate, citramalate, citraconate*, and *dimethyl citraconate*, detailing our selection rationale in Appendix A.2. Using the PubChem platform, we extracted two-dimensional *structurally similar compounds* for each. From this pool, we filtered and retained only those compounds whose canonical SMILES representations were compatible with the RDKit cheminformatics toolkit for further computational analysis. These candidate compounds remain unlabeled. With this process, we identified 567 high-potential candidate compounds, with the goal of determining the most promising ones for drug development (See Appendix A.2). A complete list of all compounds is available in our repository ^3^.

### Topological Vectors

For each compound, we extracted its topological vectors, specifically the Betti-0 vectors, using the *multipersistence framework* (Section 2). In our experiments, we combined two distinct filtration functions: *atomic weight* and *bond type*. These chemical functions were chosen intentionally as they provide an effective hierarchy among the nodes and edges, enabling meaningful decomposition into substructures. From the graph representations of each compound, by using sublevel filtrations, we derived 3 *×* 43 dimensional 2D topological vectors, which were then transformed into 1D vector representations for further analysis.

### Ranking of Compounds

We utilize the XGBoost model over the topological vectors of each compound to rank candidate compounds based on their binding affinity to the target. Positive and negative compounds are aggregated into a single dataset, which is then divided into training and validation sets using 4-fold cross-validation. To identify the best compounds, we define two ranking metrics:

- *Lowest Ranking Metric (LR):* This metric orders the compounds according to the probability for them to be positive, and then it returns the position of the positive compound that is ranked the lowest.
- *Mean Ranking Metric (MR):* Similarly to the LR, MR orders the compounds according to the probability for them to be positive, but it returns the average of the position of the positive compounds.

To verify the utility of these ranking algorithms and find optimal hyperparameters, we validate its effectivity by using their rankings of positive and negative compounds. For each ranking metric, we optimize the hyperparameter grid using both positive and negative compounds to minimize the average results obtained from a 4-fold cross-validation process. Once the optimal hyperparameter configuration is determined, we train the XGBoost model on the complete dataset, which includes both positive and negative compounds. The trained model is then applied to the candidate compound list to generate rankings based on the predicted probabilities of each compound belonging to the positive class, indicating the likelihood of binding to the target.

Since we have two metrics, two separate rankings are generated for the candidate compounds, which are then combined. Compounds with a total ranking of two are prioritized, as they exhibit a high probability of binding to the target.

### 3.1 Docking Simulations

To substantiate that identified compounds interact with AcnD specifically by involving the amino acid residues critically implicated in the active site, blind docking analysis was performed. The analysis was carried out with the molecular docking program AutoDock Vina [40,15] using the AlphaFold 3D model of AcnD that has a properly oriented catalytic 4Fe-4S cluster in its predicted active site.

AutoDock Vina is a leading docking program known for its effectiveness in predicting binding modes, particularly useful for blind docking. Blind docking enables unbiased exploration of binding sites across the entire protein surface, as it requires no prior knowledge of the binding site location. AutoDock Vina allows for searches across large defined spaces, critical for blind docking, thus enhancing flexibility to identify potential binding sites across the protein surface.

The software’s high accuracy in predicting binding modes improves the chances of identifying the true binding site, validating interactions, and revealing unexpected binding sites or modes. AutoDock Vina’s scoring functions estimate binding affinity and energy by evaluating a ligand’s orientation and position within the binding site. The Vina score, reported in kilocalories per mole (kcal/mol), quantifies binding energy—where more negative values indicate stronger interactions. This score is derived from factors like steric interactions, hydrophobic effects, and hydrogen bonding, which are weighted to produce a final binding affinity score. Compounds with lower Vina scores are ranked higher, aiding in virtual validation and prioritization for further study.

Blind docking is an efficient approach for validating lead compounds, ensuring they interact effectively with potential structural pockets before proceeding to costly experimental validations. Comparing blind docking predictions with experimental data also helps confirm that identified compounds are likely to be effective in biological systems.

Blind docking simulations are thus suitable for examining interactions between newly identified compounds and AcnD, validating their binding topologies to the active site without structural assumptions. The amino acid residues crucial for aconitase activity are highly conserved in AcnD, forming its active site, which includes positions for substrate interaction, Fe-S cluster binding, and catalytic action necessary to convert 2-methylcitrate to 2-methylisocitrate via cismethylaconitate. Like other aconitases, AcnD consists of four domains encoded by a single polypeptide [44]. Studies on pig heart mitochondrial aconitase show that residues essential for substrate binding, catalytic processing, and orientation of the 4Fe-4S cluster span all four domains: seven from domain 1, two from domain 2, seven from domain 3, and five from domain 4 [44,48]. Sequence analysis of AcnD from *Pseudomonas aeruginosa* PA01 reveals a domain arrangement like that of mitochondrial aconitase (mAcn), bacterial AcnA, and mammalian IRP [44]. The alignment shows that residues for substrate interaction, Fe-S binding, and catalysis in pig mitochondrial aconitase are similarly conserved in AcnD (Figure 7). These positions are also conserved in human mitochondrial and cytoplasmic aconitases (IRP1) (Figure 7, Table 2), supporting the idea that AcnD’s active site binds methylcitrate in a manner similar to citrate binding in mitochondrial aconitase. Although the catalytic mechanisms across aconitases are likely identical, AcnD has evolved in bacteria and fungi as a functionally analogous enzyme, targeting the intermediate methylcitrate in propionate catabolism rather than citrate from the TCA cycle. This evolution likely minimizes metabolic conflicts by enabling AcnD to convert methylcitrate to methylisocitrate independently of TCA-related aconitase activity.

**Table 1:**
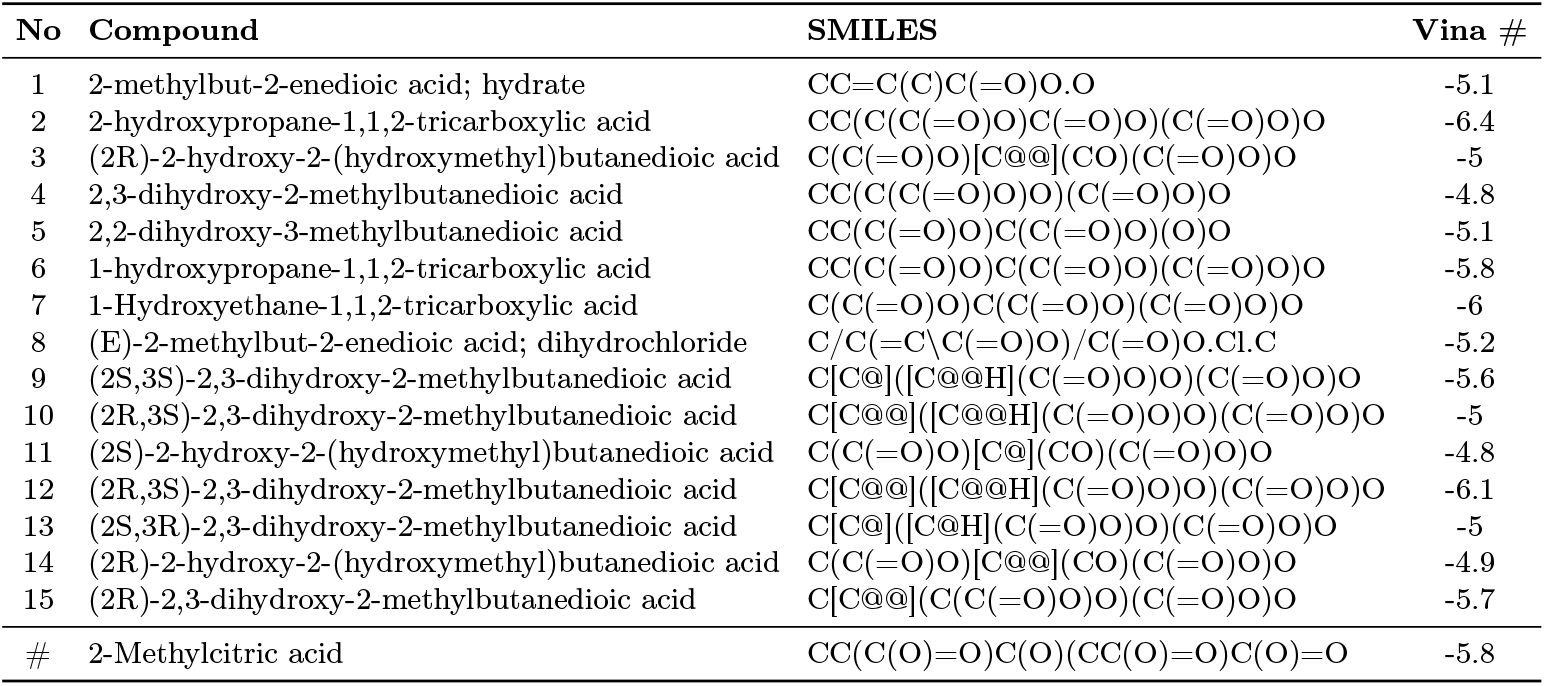
Novel compounds targeting AcnD and propionate catabolism in bacteria.

**Table 2:** Sequence Positions. Comparison of the sequence position of critical active site amino acid residues involved in substrate interaction, Fe-S cluster holding, and catalytic action between pig heart mitochondrial aconitase (ACON_PIG; NCBI Reference Sequence: NM_213954.1) and P. aeruginosa AcnD (ACND_PSEUAE; NCBI Reference Sequence: WP_003124236.1). Locations of active site amino acids on AcnD were deduced from protein sequence alignment of AcnD and functional roles were assigned in comparison to pig heart aconitase amino acids. Assignment of predicted functions is based on the numbering of amino acid residues in the multiple sequence alignment, ensuring consistency with the role of active site amino acids described for aconitase mechanism [5,48,23].

| No. | Type of Interaction | ACON_PIG | ACND_PSEUAE |
| --- | --- | --- | --- |
| 1 | Substrate interaction | Gln114 | Gln103 |
| 2 | Catalytic | Asp127 | Asp119 |
| 3 | Substrate interaction | His128 | His120 |
| 4 | Catalytic | His174 | His173 |
| 5 | Substrate interaction | Asp192 | Asp200 |
| 6 | Substrate interaction | Ser193 | Ser201 |
| 7 | Catalytic | His194 | His202 |
| 8 | Substrate interaction | Asn285 | Asn293 |
| 9 | Catalytic | Glu289 | Glu297 |
| 10 | Fe-S cluster interaction | Cys385 | Cys411 |
| 11 | Fe-S cluster interaction | Cys448 | Cys477 |
| 12 | Fe-S cluster interaction | Cys451 | Cys480 |
| 13 | Substrate interaction | Ile452 | Asn481 |
| 14 | Substrate interaction | Gly453 | Gly482 |
| 15 | Substrate interaction | Asn473 | Asn509 |
| 16 | Substrate interaction | Arg474 | Arg510 |
| 17 | Substrate interaction | Arg479 | Arg515 |
| 18 | Substrate interaction | Arg534 | Arg574 |
| 19 | Catalytic | Ser669 | Ser751 |
| 20 | Substrate interaction | Ser670 | Ser752 |
| 21 | Substrate interaction | Arg671 | Arg753 |

The docking simulation results confirmed that the newly identified compounds can effectively interact with the AcnD active site (Figures 8 to 10). These simulations revealed significant interactions within the active site of AcnD, involving amino acid residues crucial for binding and processing the natural ligand, methylcitrate. The confirmation of these contact sites supports the compounds’ structural alignment with AcnD’s native substrate (2-methylcitrate), product (2-isomethylcitrate), and the reaction intermediate (cis-methylaconitate). For instance, *2,3-dihydroxy-2-methylbutanedioic acid* exhibited interactions within AcnD’s active site, following a predictable binding pattern with key residues (Figure 2). Overall, the docking simulations validated both the molecular structure and potential binding efficacy of these compounds within AcnD’s active site. The compounds maintained their conformational stability and fit well within the AcnD pocket, underscoring their capacity to engage key residues essential for effective ligand positioning. This highlights the compounds’ potential to compete with methylcitrate, thereby inhibiting AcnD activity. Hence, these compounds represent promising drug leads for targeting the methylcitrate pathway, warranting further experimental investigation into potential therapeutic applications.

## 4 Conclusion

This study demonstrates how the integration of topology modeling, machine learning (ML), and molecular docking can enhance ligand-based drug discovery. Topology modeling provides a structural framework for molecular features, generating topology vectors that enable ML to prioritize and search vast compound datasets. Topological descriptors are critical in elucidating structure-activity relationships, with topologically categorized molecules sharing essential substructures that favor specific and stronger interactions within target sites. Docking simulations bolster the predictive reliability of topological ranking by detailing interactions and illustrating how common substructures fit within target binding sites. Empirical binding site validation and interaction pattern confirmation enable prioritization of compounds with favorable interactions, improving their efficacy potential. By considering the geometric and energetic fit alongside interaction profiles, predictions become more refined, aiding the design of new compounds with optimized properties. High-ranking topologies serve as a robust foundation for lead optimization by representing plausible molecular scaffolds suitable for targeted modifications. This process narrows the chemical space to structurally similar compounds with beneficial interaction profiles, facilitating efficient lead compound enhancement and optimizing drug-like properties. Prioritizing high-potential compounds early accelerates discovery, improves accuracy in predicting compound-target interactions, and reduces time, cost, and failure risk in later drug development stages, saving significant resources.

Topologically validated compounds that confirm drug-target interactions offer valuable scaffolds for chemical modifications, enhancing specificity and efficacy. Enzyme-validated topologies are particularly useful for targeting metabolic reactions, making substrate-mimicking compounds strong inhibitor candidates. Future work will expand this approach to other metabolic pathways, using topologies designed to mimic various substrates and intermediates, potentially disrupting microbial pathways beyond the methylcitrate pathway. This could lead to antimicrobial agents for resistant bacteria. Combining topology modeling with AI techniques, such as generative models, may further improve scaffold specificity and efficacy, promising advancements in metabolic disease treatments and other therapeutic areas.

## Appendix

### A Further Details

#### A.1 Multifiltrations and Topological Vectors

In this part, we give an explicit toy example of our topological vectors using multiparameter persistence (MP). In Figure 5, we give the molecular graph for *methyl (E)-2-methylhex-2-en-4-ynoate*. In Figure 6, we give a toy illustration for its multifiltration for three thresholds for bond strength (1,2,3) and three thresholds for atomic number (1,6,8). By using we obtain two 3 × 3 vectors 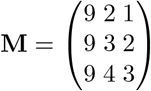, where each entry represent the number of components in corresponding cells in Figure 6.

**Fig. 5:**
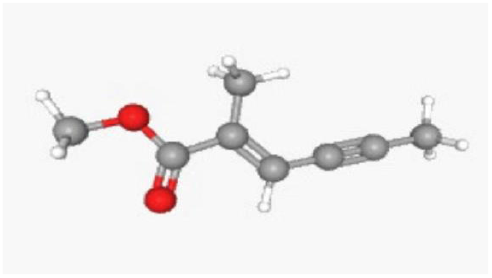
Sample compound.

**Fig. 6:**
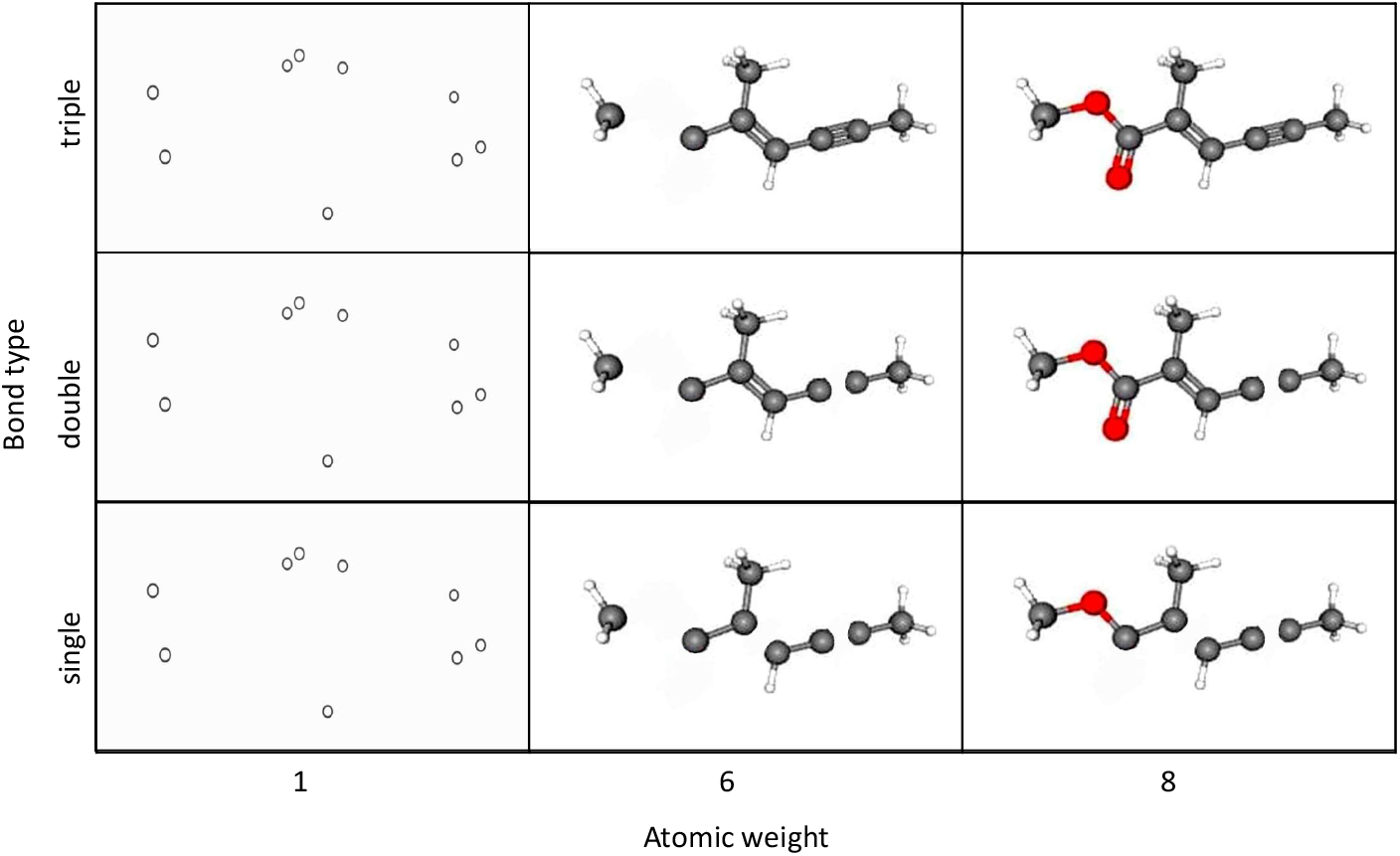
Toy Example for Multifiltration. In the 3 *×* 3 grid above, we give a toy example for a multifiltration. *x*-axis represents the 3 thresholds (1, 6, 8) for atomic numbers, while *y*-axis represents 3 thresholds (1,2,3) for bond strength. We activate each node in the corresponding cell if its atomic number reaches the corresponding threshold. We activate each edge if the bond strength reaches to corresponding cell. For example, the middle cell represents all nodes whose atomic number is ≤ 6 and all edges whose strength ≤ 2.

#### A.2 Creation of Compound Library

In this study, we applied a topological machine learning model to identify new compounds that target AcnD, an enzyme essential in microbial propionate catabolism within the methylcitrate cycle. AcnD, an aconitase-like enzyme, catalyzes the conversion of methylcitrate to methylisocitrate—an essential reaction in this metabolic cycle. AcnD shares conserved amino acid sequences and domain architecture with other aconitase family proteins, such as AcnA (part of the TCA cycle) and IRP (cytosolic aconitase). This structural conservation suggests a similar active site structure and catalytic mechanism, leading to selective recognition and binding of methylcitrate as its substrate.

Methylcitrate, specifically *2-methylcitrate*, shares topological features with compounds such as *citrate, isocitrate, 2-isomethylcitrate, aconitate*, and *methylaconitate*, which all feature a core tricarboxylic acid structure central to their function. This core includes three carboxyl groups attached to a carbon backbone, facilitating binding and catalysis by enzymes like AcnD. The presence of functional groups like hydroxyls and carboxyls is crucial for biochemical interactions, enabling hydrogen bonding and ionic interactions within the enzyme’s active site. The isomerization capacity of these compounds, particularly with the addition of a methyl group in 2-methylcitrate and its isomers, introduces slight topological variations that do not alter the overall molecular structure significantly. This shared topology among these compounds supports their compatibility with the aconitase family’s active sites, including AcnD.

These shared features enabled us to define a topological representation that captures essential aspects of 2-methylcitrate’s molecular shape, size, and functional groups. This topology vector, derived from relevant molecular features, serves as an input for machine learning algorithms to search for similar topological patterns in large compound libraries, facilitating the identification of structurally related compounds with potential bioactivity toward AcnD.

### B Docking Simulations

Molecular docking simulations (Figures 8 to 10) were conducted using AutoDock Vina version 1.1.2 [40,15] to explore binding interactions with the Fe-S-dependent enzyme AcnD from *Pseudomonas aeruginosa* (UniProt A0A8B4ZGT7). The protein’s predicted three-dimensional structure, featuring an Fe-S cluster, was retrieved from the AlphaFold Protein Structure Database. Preparation of the protein for docking involved removing water molecules and optimizing protonation states to match physiological pH (7.4). Hydrogen atoms were added, and the structure was processed in AutoDock Tools (ADT) version 1.5.7 to parameterize the Fe-S cluster. Polar hydrogens were added, and Gasteiger charges assigned, after which non-polar hydrogens were merged, saving the prepared structure as AcnD_Model_Fe4S4.pdbqt.

Ligands were prepared by converting their structures from SDF to PDB format with Open Babel version 3.1.1, followed by energy minimization using the MMFF94 force field to achieve stable conformations. After adding hydrogen atoms and assigning partial charges using Gasteiger methods, each ligand was loaded into ADT, where Gasteiger charges were recalculated, and rotatable bonds defined automatically. The prepared ligands were saved individually as ligand.pdbqt files.

Docking simulations were set up with a grid box defined from fpocket version 3.0 analysis of potential binding sites. Using default parameters, fpocket ranked predicted cavities; the top cavity, representing the active site with the Fe4S4 cluster, was selected. Center coordinates (x=7, y=2, z=-1) and dimensions (x=31, y=35, z=35) of this pocket were obtained from the pockets.pdb file. Docking was performed with the exhaustiveness parameter set to 8, balancing computational efficiency with thorough search depth.

Docking scores (Vina scores) reflect binding affinities and are derived from weighted interactions, including steric (van der Waals) forces, hydrophobic clustering, hydrogen bonding, and entropic contributions from rotational and translational constraints. Scores account for interactions contributing most significantly to binding energy among atoms involved. The amino acids defining the docking pocket and their positions on AcnD are listed for each compound, with residues critical to the active site highlighted in red. Close-up views of binding pockets and corresponding 2D interaction diagrams illustrate detailed ligand-protein interactions, supported by SMILES codes for compound structure translation.

#### B.1 Protein Sequence Alignments

In Figure 7, we give protein sequence alignments of pig mitochondrial aconitase (ACON_PIG; NCBI Reference Sequence: NM_213954.1), human mitochondrial aconitase (ACOC_HUMAN_Mitochondrial; UniProtKB/Swiss-Prot: Q99798.2), human cytoplasmic aconitase (IRP1) (ACOC_HUMAN_Cytoplasmic; NCBI Reference Sequence: NP_001265281.1), and Pseudomonas aeruginosa AcnD (ACND_PSEUAE; NCBI Reference Sequence: WP_003124236.1). Conserved residues are indicated with * in the consensus. Critical amino acid residues that are attributed to substrate interaction, Fe-S cluster binding, and catalytic action in the active site of pig mitochondrial aconitase and the putative locations of the corresponding conserved amino acids residues in AcnD are highlighted.

**Fig. 7:**
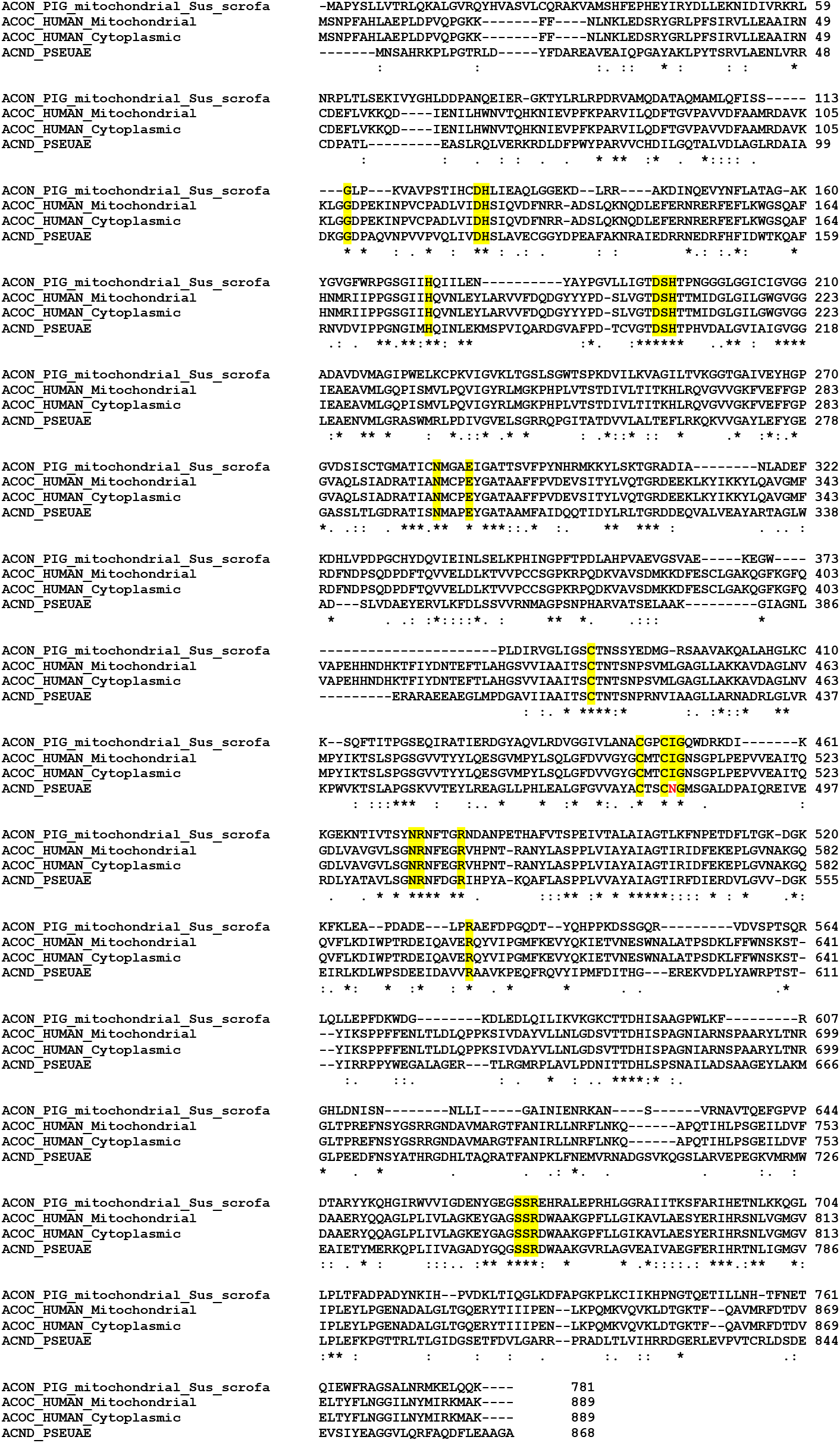
Sequence Alignments.

#### B.2 Results of Blind Docking Analysis

In Figures 8 to 10, we present our molecular docking simulations conducted using AutoDock Vina version 1.1.2 [40,15]. We used the three-dimensional structure of Pseudomonas aeruginosa AcnD (Fe/S-dependent 2-methylisocitrate dehydratase; UniProt A0A8B4ZGT7), which includes the 4Fe4S cluster, for docking studies. The protein’s predicted structure was obtained from the AlphaFold Protein Structure Database. For docking preparation, water molecules were removed from the AcnD structure, and protonation states were adjusted to physiological pH (7.4). Hydrogen atoms were added, and the structure was loaded into AutoDock Tools (ADT) version 1.5.7 to parameterize the 4Fe4S cluster using AutoDock metal parameters, followed by the addition of polar hydrogens and assignment of Gasteiger charges. Non-polar hydrogens were merged, and the prepared protein was saved as AcnD_Model_Fe4S4.pdbqt.

For ligand preparation, Open Babel version 3.1.1 was used to convert compound structures from SDF to PDB format. Each ligand was energy-minimized using the MMFF94 force field to achieve stable conformations, then loaded individually into ADT with added hydrogens and assigned Gasteiger charges. Rotatable bonds were defined automatically, and each ligand was saved as a separate ligand.pdbqt file.

Docking was performed individually for each ligand using grid box parameters obtained from fpocket analysis, with the exhaustiveness parameter set to 8 for an optimal balance between computational efficiency and thoroughness. Binding sites on AcnD were identified using fpocket version 3.0, with predicted cavities ranked by potential binding suitability. The top-ranking cavity, corresponding to the AcnD active site containing the Fe4S4 cluster, was selected for docking. The pocket’s coordinates (centered at x=7, y=2, z=-1) and dimensions (x=31, y=35, z=35) were extracted from fpocket output.

Docking scores (Vina scores) reflect estimated binding affinities of the compounds to AcnD, based on an empirical model that incorporates steric, hydrophobic, hydrogen bonding, and entropy-driven interactions. These scores provide insight into the relative binding strengths of different compounds. Each docking pocket’s amino acids and sequence positions on AcnD are listed, with red font used to indicate amino acids implicated in the active site. Close-up 3D views of the binding pockets and accompanying 2D interaction diagrams illustrate the spatial position, orientation, and specific amino acid interactions within AcnD’s active site.

**Fig. 8:**
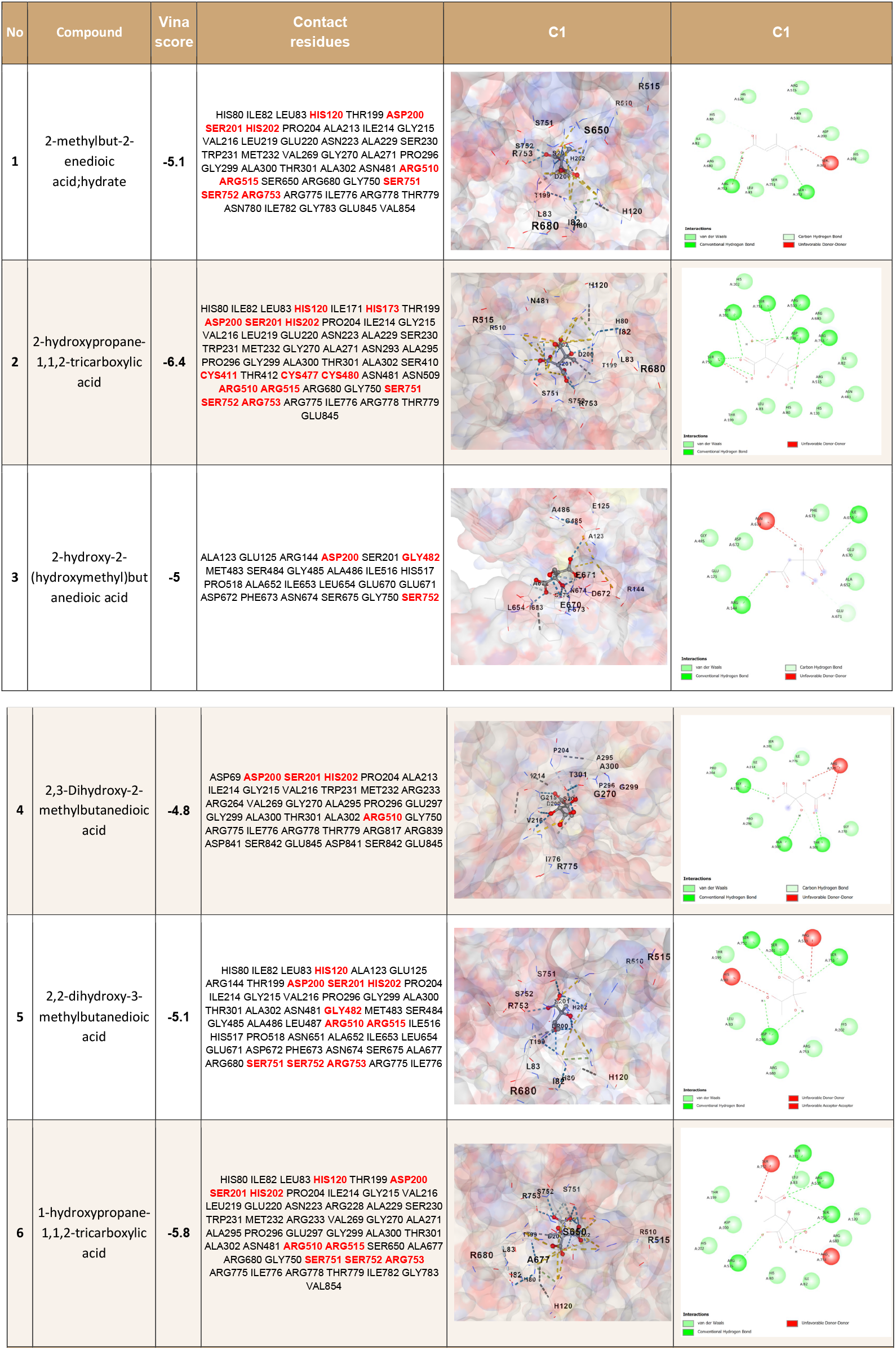
Docking simulations.

**Fig. 9:**
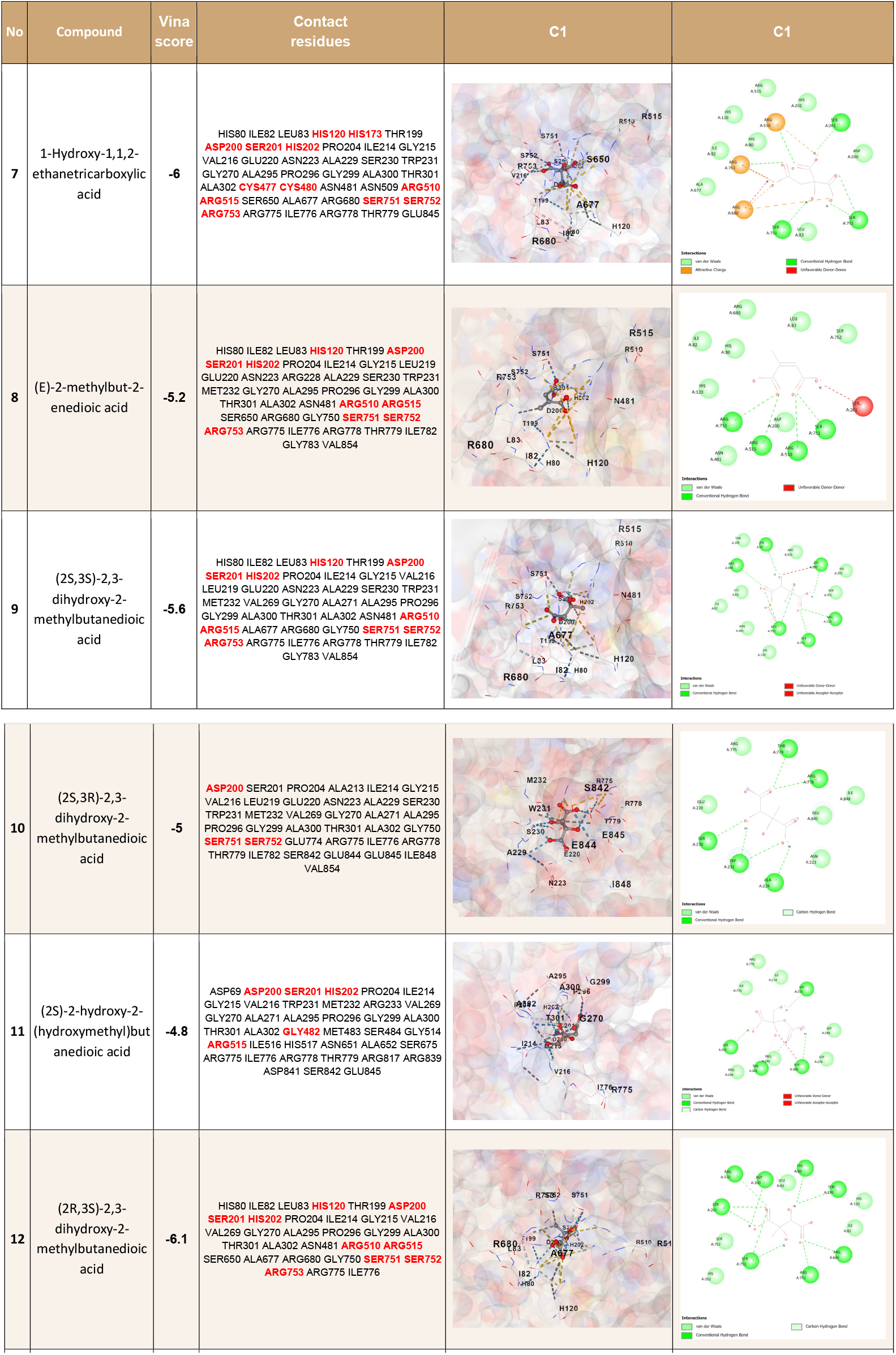
Docking simulations.

**Fig. 10:**
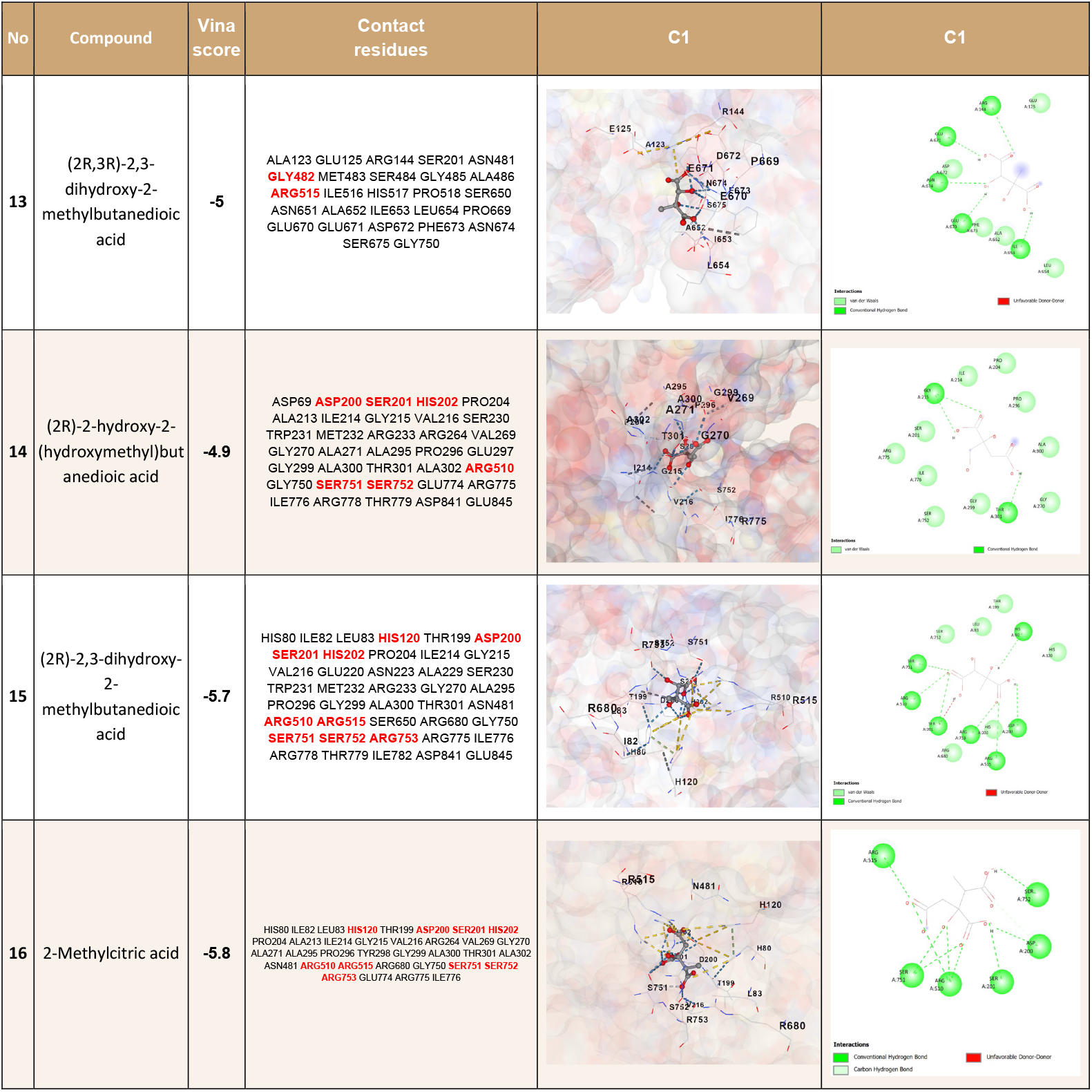
Docking simulations.

## Footnotes

3 https://github.com/AstritTola/Molecular-Compounds-Targeting

## References

1. Agüero-Chapin, G., Antunes, A., Marrero-Ponce, Y.: A 2022 update on computational approaches to the discovery and design of antimicrobial peptides. Antibiotics 12(6), 1011 (2023). 10.3390/antibiotics12061011

2. Aktas, M.E., Akbas, E., El Fatmaoui, A.: Persistence homology of networks: methods and applications. Applied Network Science 4(1), 1–28 (2019)

3. Ali, D., Asaad, A., Jimenez, M.J., Nanda, V., Paluzo-Hidalgo, E., Soriano-Trigueros, M.: A survey of vectorization methods in topological data analysis. IEEE Transactions on Pattern Analysis and Machine Intelligence (2023)

4. Aronica, P., Reid, L., Desai, N., et al.: Computational methods and tools in antimicrobial peptide research. Journal of Chemical Information and Modeling 61(7), 3172–3196 (Jul 2021). 10.1021/acs.jcim.1c00175

5. Beinert, H., Kennedy, M.: Aconitase, a two-faced protein: enzyme and iron regulatory factor. FASEB J. 7(15), 1442–1449 (Dec 1993). 10.1096/fasebj.7.15.8262329

6. Brock, M., Maerker, C., Schütz, A., Völker, U., Buckel, W.: Oxidation of propionate to pyruvate in escherichia coli. involvement of methylcitrate dehydratase and aconitase. European Journal of Biochemistry 269(24), 6184–6194 (Dec 2002). 10.1046/j.1432-1033.2002.03336.x

7. Brock, M., Fischer, R., Linder, D., Buckel, W.: Methylcitrate synthase from aspergillus nidulans: implications for propionate as an antifungal agent. Molecular Microbiology: Original Articles 35(5), 961–973 (2000)

8. Chalmers, R., Lawson, A., Chalmers, R., Lawson, A.: Disorders of propionate and methylmalonate metabolism. Organic Acids in Man: Analytical Chemistry, Biochemistry and Diagnosis of the Organic Acidurias pp. 296–331 (1982)

9. Coskunuzer, B., Akçora, C.G.: Topological methods in machine learning: A tutorial for practitioners. arXiv preprint arXiv:2409.02901 (2024)

10. Cui, G., Zhang, Y., Xu, X., Liu, Y., Li, Z., Wu, M., Liu, J., Gan, J., Liang, H.: Pmir senses 2-methylisocitrate levels to regulate bacterial virulence in pseudomonas aeruginosa. Science Advances 8(49), eadd4220 (2022)

11. Dadgostar, P.: Antimicrobial resistance: implications and costs. Infection and drug resistance pp. 3903–3910 (2019)

12. Demir, A., Coskunuzer, B., Gel, Y., Segovia-Dominguez, I., Chen, Y., Kiziltan, B.: Todd: Topological compound fingerprinting in computer-aided drug discovery. Advances in Neural Information Processing Systems 35, 27978–27993 (2022)

13. Dey, T.K., Wang, Y.: Computational Topology for Data Analysis. Cambridge University Press (2022)

14. Dolan, S.K., Wijaya, A., Geddis, S.M., Spring, D.R., Silva-Rocha, R., Welch, M.: Loving the poison: the methylcitrate cycle and bacterial pathogenesis. Microbiology 164(3), 251–259 (2018)

15. Eberhardt, J., Santos-Martins, D., Tillack, A.F., Forli, S.: Autodock vina 1.2.0: New docking methods, expanded force field, and python bindings. Journal of Chemical Information and Modeling (2021)

16. Garvey, G.S., Rocco, C.J., Escalante-Semerena, J.C., Rayment, I.: The three-dimensional crystal structure of the prpf protein of shewanella oneidensis complexed with trans-aconitate: Insights into its biological function. Protein science 16(7), 1274–1284 (2007)

17. Grimek, T.L., Escalante-Semerena, J.C.: The acnd genes of shewenella oneidensis and vibrio cholerae encode a new fe/s-dependent 2-methylcitrate dehydratase enzyme that requires prpf function in vivo. Journal of bacteriology 186(2), 454–462 (2004)

18. Hammelman, T., O’Toole, G., Trzebiatowski, J., Tsang, A., Rank, D., Escalante-Semerena, J.: Identification of a new prp locus required for propionate catabolism in salmonella typhimurium lt2. FEMS microbiology letters 137(2-3), 233–239 (1996)

19. Hatcher, A.: Algebraic Topology. Cambridge University Press (2002)

20. Horswill, A.R., Escalante-Semerena, J.C.: Propionate catabolism in salmonella typhimurium lt2: two divergently transcribed units comprise the prp locus at 8.5 centisomes, prpr encodes a member of the sigma-54 family of activators, and the prpbcde genes constitute an operon. Journal of bacteriology 179(3), 928–940 (1997)

21. Jaiswal, A., Babu, A.R., Zadeh, M.Z., Banerjee, D., Makedon, F.: A survey on contrastive self-supervised learning. Technologies 9(1), 2 (2020)

22. Khan, S., Madhi, S., Olwagen, C.: Structure-based identification of novel inhibitors targeting the enoyl-acp reductase enzyme of acinetobacter baumannii. Scientific Reports 13, 21331 (2023). 10.1038/s41598-023-48696-z

23. Lloyd, S., Lauble, H., Prasad, G., Stout, C.: The mechanism of aconitase: 1.8 a resolution crystal structure of the s642a:citrate complex. Protein Sci. 8(12), 2655–2662 (Dec 1999). 10.1110/ps.8.12.2655

24. Loiseaux, D., Scoccola, L., Carrière, M., Botnan, M.B., Oudot, S.: Stable vectorization of multiparameter persistent homology using signed barcodes as measures. Advances in Neural Information Processing Systems 36 (2024)

25. London, R.E., Allen, D.L., Gabel, S.A., DeRose, E.F.: Carbon-13 nuclear magnetic resonance study of metabolism of propionate by escherichia coli. Journal of bacteriology 181(11), 3562–3570 (1999)

26. Melo, M., Maasch, J., de la Fuente-Nunez, C.: Accelerating antibiotic discovery through artificial intelligence. Communications Biology 4(1), 1050 (Sep 2021). 10.1038/s42003-021-02586-0

27. Meng, Q., et al.: Dmamp: A deep-learning model for detecting antimicrobial peptides and their multi-activities. IEEE/ACM Transactions on Computational Biology and Bioinformatics (2024). 10.1109/TCBB.2024.3439541

28. Meng, Z., Anand, D.V., Lu, Y., Wu, J., Xia, K.: Weighted persistent homology for biomolecular data analysis. Scientific reports 10(1), 2079 (2020)

29. Mukherjee, S., Samaga, S.N., Xin, C., Oudot, S., Dey, T.K.: D-gril: End-to-end topological learning with 2-parameter persistence. arXiv preprint arXiv:2406.07100 (2024)

30. Murray, C.J., Ikuta, K.S., Sharara, F., Swetschinski, L., Aguilar, G.R., Gray, A., Han, C., Bisignano, C., Rao, P., Wool, E., et al.: Global burden of bacterial antimicrobial resistance in 2019: a systematic analysis. The lancet 399(10325), 629–655 (2022)

31. Nguyen, L., Garcia, J., Gruenberg, K., MacDougall, C.: Multidrug-resistant pseudomonas infections: hard to treat, but hope on the horizon? Current infectious disease reports 20, 1–10 (2018)

32. Prestinaci, F., Pezzotti, P., Pantosti, A.: Antimicrobial resistance: a global multi-faceted phenomenon. Pathogens and global health 109(7), 309–318 (2015)

33. Qin, S., Xiao, W., Zhou, C., Pu, Q., Deng, X., Lan, L., Liang, H., Song, X., Wu, M.: Pseudomonas aeruginosa: pathogenesis, virulence factors, antibiotic resistance, interaction with host, technology advances and emerging therapeutics. Signal Transduction and Targeted Therapy 7(1), 199 (Jun 2022). 10.1038/s41392-022-01056-1

34. Rocco, C.J., Wetterhorn, K.M., Garvey, G.S., Rayment, I., Escalante-Semerena, J.C.: The prpf protein of shewanella oneidensis mr-1 catalyzes the isomerization of 2-methyl-cis-aconitate during the catabolism of propionate via the acnd-dependent 2-methylcitric acid cycle. Plos one 12(11), e0188130 (2017)

35. Sabe, V.T., Ntombela, T., Jhamba, L.A., Maguire, G.E., Govender, T., Naicker, T., Kruger, H.G.: Current trends in computer aided drug design and a highlight of drugs discovered via computational techniques: A review. European Journal of Medicinal Chemistry 224, 113705 (2021)

36. Sadybekov, A.V., Katritch, V.: Computational approaches streamlining drug discovery. Nature 616(7958), 673–685 (2023)

37. Shimizu, Y., Yonezawa, T., Sakamoto, J., et al.: Identification of novel inhibitors of keap1/nrf2 by a promising method combining protein–protein interaction-oriented library and machine learning. Scientific Reports 11, 7420 (2021). 10.1038/s41598-021-86616-1

38. Suvorova, I., Ravcheev, D., Gelfand, M.: Regulation and evolution of malonate and propionate catabolism in proteobacteria. Journal of bacteriology 194(12), 3234–3240 (2012)

39. The World Bank: Drug-resistant infections: A threat to our economic future. https://www.worldbank.org/en/topic/health/publication/drug-resistant-infections-a-threat-to-our-economic-future (2017)

40. Trott, O., Olson, A.J.: Autodock vina: improving the speed and accuracy of docking with a new scoring function, efficient optimization, and multithreading. Journal of Computational Chemistry 31(2), 455–461 (2010)

41. Upton, A.M., McKinney, J.D.: Role of the methylcitrate cycle in propionate metabolism and detoxification in mycobacterium smegmatis. Microbiology 153(12), 3973–3982 (2007)

42. Verma, Y., Souza, A.H., Garg, V.: Topological neural networks go persistent, equivariant, and continuous. arXiv preprint arXiv:2406.03164 (2024)

43. Wan, F., Wong, F., Collins, J., et al.: Machine learning for antimicrobial peptide identification and design. Nature Reviews Bioengineering 2, 392–407 (2024). 10.1038/s44222-024-00152-x

44. Watanabe, S., Murase, Y., Watanabe, Y., Sakurai, Y., Tajima, K.: Crystal structures of aconitase x enzymes from bacteria and archaea provide insights into the molecular evolution of the aconitase superfamily. Communications Biology 4(1), 687 (2021)

45. Wever, M.J., Scommegna, F.R., Egea-Rodriguez, S., et al.: Structure-based discovery of first inhibitors targeting the helicase activity of human pif1. Nucleic Acids Research (2024). 10.1093/nar/gkae897

46. Wijaya, A.J.: Characterization and targeting of the 2-methylcitrate cycle in Pseudomonas aeruginosa. Phd thesis, University of Cambridge (2021), 10.17863/CAM.64463

47. World Health Organization: Antimicrobial resistance. https://www.who.int/health-topics/antimicrobial-resistance (2024)

48. Zheng, L., Kennedy, M.C., Beinert, H., Zalkin, H.: Mutational analysis of active site residues in pig heart aconitase. Journal of Biological Chemistry 267(11), 7895–7903 (1992)

